# Application of the anatomical fiducials framework to a clinical dataset of patients with Parkinson’s disease

**DOI:** 10.1101/2020.12.17.423333

**Authors:** Mohamad Abbass, Greydon Gilmore, Alaa Taha, Ryan Chevalier, Magdalena Jach, Terry M. Peters, Ali R. Khan, Jonathan C. Lau

**Affiliations:** Department of Clinical Neurological Sciences, London Health Sciences Centre, Western University, London, ON, Canada; School of Biomedical Engineering, Western University, London, Ontario, Canada; Department of Physiology, Western University, London, Ontario, Canada; Imaging Research Laboratories, Robarts Research Institute, Western University, London, Ontario, Canada; Centre for Functional and Metabolic Mapping, Robarts Research Institute, Western University, London, Ontario, Canada; Department of Medical Biophysics, Western University, London, Ontario, Canada; Brain and Mind Institute, Western University, London, Ontario, Canada; Graduate Program in Neuroscience, Western University, London, Ontario, Canada; Department of Neurosurgery, Emory University, Atlanta, Georgia, USA

**Keywords:** Registration, Accuracy, Fiducials, Deep brain stimulation, Parkinson’s Disease, Biomarker

## Abstract

Establishing spatial correspondence between subject and template images is necessary in neuroimaging research and clinical applications such as brain mapping and stereotactic neurosurgery. Our anatomical fiducials (AFIDs) framework has recently been validated to serve as a quantitative measure of image registration based on salient anatomical features. In this study, we sought to apply the AFIDs protocol to the clinic, focusing on structural magnetic resonance images obtained from patients with Parkinson’s Disease (PD). We confirmed AFIDs could be placed to millimetric accuracy in the PD dataset with results comparable to those in normal control subjects. We evaluated subject-to-template registration using this framework by aligning the clinical scans to standard template space using a robust open preprocessing workflow. We found that registration errors measured using AFIDs were higher than previously reported, suggesting the need for optimization of image processing pipelines for clinical grade datasets. Finally, we examined the utility of using point-to-point distances between AFIDs as a morphometric biomarker of PD, finding evidence of reduced distances between AFIDs that circumscribe regions known to be affected in PD including the substantia nigra. Overall, we provide evidence that AFIDs can be successfully applied in a clinical setting and utilized to provide localized and quantitative measures of registration error. AFIDs provide clinicians and researchers with a common, open framework for quality control and validation of spatial correspondence and the location of anatomical structures, facilitating aggregation of imaging datasets and comparisons between various neurological conditions.

## Introduction

Non-invasive imaging techniques such as magnetic resonance imaging (MRI) have allowed for insights into the anatomy and function of the central nervous system. A critical aspect in neuroimaging research and clinical application is to establish accurate spatial correspondence between images (Chakravarty et al. 2008), allowing for the combination and comparison of multimodal data across subjects and populations. Establishing spatial correspondence requires the specification of a common stereotactic 3D coordinate reference frame, and the registration of 3D images to that reference frame (Pirker and Katzenschlager 2017). Researchers have established numerous common reference frames, including those based on both individuals and populations (Fonov et al. 2011).

Establishing correspondence between various brain images has historically relied on linear transformations (Fonov et al. 2011; Evans et al. 1992). Over the last few decades, various nonlinear transformations have been implemented allowing for more accurate registration between brain images (Fonov et al. 2011). These transformations have most commonly been evaluated using measures of overlap between regions of interest (ROIs). ROIs that have been used include subcortical structures such as the thalamus or areas of the basal ganglia (Lau et al. 2019; Poldrack 2007). Measures of spatial correspondence within these relatively large ROIs are known to be quite coarse and fail to capture subtle misregistration between images (Rohlfing 2012; Lau et al. 2019). Inspired by classical stereotactic methods (Talairach et al. 1958), a set of anatomical fiducials (AFIDs) were validated using an open framework and proposed as an intuitive way to quantify alignment using point-based distance measures between brain structures. This method was validated in individual subject and template scans and was found to be more sensitive to registration errors than ROI-based voxel overlap measures (Lau et al. 2019).

With the increasing use of MRI in research and clinical settings, accurate assessment of registration between image sequences is necessary. Since clinical outcomes in stereotactic neurosurgery depend on accuracy at the millimeter scale (Li et al. 2016), a robust framework for assessing correspondence between brain images is required for optimal neurosurgical planning. In this work, we sought to evaluate the reproducibility and utility of the AFIDs framework in a clinical population with Parkinson’s Disease (PD).

## Material and Methods

All raw and processed data along with the processing scripts that were used in this manuscript are available at: https://github.com/afids/afids-clinical. This repository is licensed under the MIT License.

### Subject demographics and MRI acquisition

Subject scans used in this study were obtained from 39 individuals diagnosed with PD (age: 60.2 ± 6.8, sex: 33.3% female). For all subjects, the MRI sequence used was a post gadolinium enhanced volumetric T1-weighted (T1w) image (echo time = 1.5 ms, inversion time = 300 ms, flip angle = 20°, receiver bandwidth = 22.73 kHz, field of view = 26 cm x 26 cm, matrix size = 256 × 256, slice thickness = 1.4 mm, resolution = 1.25 × 1.25 × 1.50 mm) (Signa, 1.5 T, General Electric, Milwaukee, Wisconsin, USA). The subject data were collected at University Hospital in London, ON, Canada. The study was approved by the Human Subject Research Ethics Board (HSREB) office at the University of Western Ontario (REB# 109045).

### AFID placement

The individual scans were imported into 3D Slicer version 4.10.0 (Fedorov et al. 2012). The subject scans were first transformed into anterior commissure (AC)-posterior commissure (PC) space (AC-PC space), and the raters were required to initially place 4 of the AFIDs, which included: AC (AFID01), PC (AFID02) and two additional on the midline. The built-in “AC-PC Transform” function in 3D Slicer was used to align the AC-PC horizontally in-line in the anteroposterior plane. Adequate alignment was subjectively judged by each rater, who then placed the remaining AFIDs as previously outlined (Lau et al. 2019). An interactive 3-dimensional schematic brain with all AFIDs labelled can be found in the supplementary material (Online Resource 1) for reference.

Five raters were initially trained to place AFIDs using publicly available brain images: MNI152NLin2009bAsym (Fonov et al. 2011; Ciric et al. 2021), deepbrain7t (Lau et al. 2017) and PD25-T1MPRAGE (Xiao et al. 2017). Each template has a set of ideal AFID coordinates (ground truth), which represents the mean AFID coordinate between a set of experienced raters. The ground truth standards are included in the GitHub repository (https://github.com/greydongilmore/afids-clinical/data/fid_standards). Quality assurance was performed to ensure each rater was placing the AFIDs on the templates below a minimum threshold of error (Euclidean error < 2.00 mm when compared with ground truth placements). Once the raters had received adequate feedback about their initial ratings during the training phase, they then independently performed the AFIDs protocol in the subject scans. Two raters (MA and GG) had prior neuroanatomy experience and were deemed “expert”, while three (AT, MJ and RC) had no prior neuroanatomy experience and were deemed “novice”. The novice raters had no experience with medical imaging so additional training was provided on navigating an MRI sequence in 3D Slicer (i.e. left/right, axial/coronal/sagittal views etc.). A total of 6240 AFIDs were placed.

### Analysis in subject space

The 3D coordinates of each AFID were exported and subsequently analyzed in MATLAB (vR2018b). The anatomical fiducial localization error (AFLE) was calculated as the Euclidean distance between each individually placed AFID and the group mean, in each of the 32 AFIDs in each scan. Therefore, 6240 AFLE measurements were made for each manually placed AFID. Outliers were determined as having an AFLE of greater than 10.0 mm and are reported in the results.

To determine each rater’s deviation from the group mean, the mean rater AFLE across all 39 subjects was calculated for each AFID. AFLE was then dichotomized between expert and novice raters by calculating the mean AFLE among these two groups. Wilcoxon rank-sum tests were used to determine significance in AFLEs between expert and novice raters. Bonferroni correction was used to account for multiple comparisons with an adjusted p-value of 0.05/32 as a threshold for significance. The overall AFLE for each AFID was then calculated as the mean AFLE across all raters.

Rater reliability was assessed using intraclass correlation (ICC), which was calculated in each dimension. A two-way random effects model with single measurement type was used, ICC(2,1) as determined by Shrout and Fliess (Shrout and Fleiss 1979). ICC among all raters, expert raters and novice raters was calculated.

### Analysis in MNI Space

To assess and quantify registration error, the subject scans were non-linearly transformed to MNI152NLin2009cAsym brain template space using fMRIPrep 1.5.4 ((Esteban et al. 2019); RRID:SCR_016216), which is based on Nipype 1.3.1 ((Gorgolewski et al. 2011); RRID:SCR_002502). Specifically, the T1-weighted (T1w) image was corrected for intensity nonuniformity (INU) with N4BiasFieldCorrection (Tustison et al. 2010), distributed with ANTs 2.2.0 ((Avants et al. 2008); RRID:SCR_004757), and used as T1w-reference throughout the workflow. The T1w reference was then skull-stripped with a Nipype implementation of the antsBrainExtraction.sh workflow (from ANTs), using OASIS30ANTs as the target template. Brain tissue segmentation of cerebrospinal fluid, white-matter and gray-matter was performed on the brain-extracted T1w using the fast algorithm from FSL 5.0.9 ((Zhang et al. 2001); RRID:SCR_002823). Volume-based spatial normalization to one standard space (MNI152NLin2009cAsym) was performed using a symmetric diffeomorphic image registration method (antsRegistration; ANTs 2.2.0), using brain-extracted versions of both T1w reference and the T1w template. The following template was selected for spatial normalization: ICBM 152 Nonlinear Asymmetrical template version 2009c ((Fonov et al. 2009); RRID:SCR_008796; TemplateFlow ID: MNI152NLin2009cAsym). Many internal operations of fMRIPrep use Nilearn 0.6.0 ((Abraham et al. 2014); RRID:SCR_001362), mostly within the functional processing workflow. For more details of the pipeline, see the section corresponding to workflows in fMRIPrep’s documentation.

We transformed each individually placed AFID to MNI space, and the mean coordinates of each AFID across all raters to MNI space. We calculate the Euclidean distance between each individually placed AFID transformed to MNI space and the group mean for each AFID placed in MNI space. We term this the *real-world* Anatomical Fiducial Registration Error (AFRE). The mean real-world AFRE across all subjects and raters was then calculated in the same manner as for the AFLE. We then calculate the Euclidean distance from the mean AFID transformed to MNI space, obtained by averaging the coordinates across all raters, and termed this the *consensus* AFRE, consistent with our definition in the original manuscript (Lau et al. 2019). The real-world AFRE represents the expected AFRE obtained by a single rater, and we focussed on this analysis since it most represents the situation in a clinical setting, although we also computed the consensus AFRE since it represents a better overall measure of registration error within our clinical sample and is directly comparable to our prior work. A schematic illustrating these measures is presented in Fig.1.

**Fig. 1.**
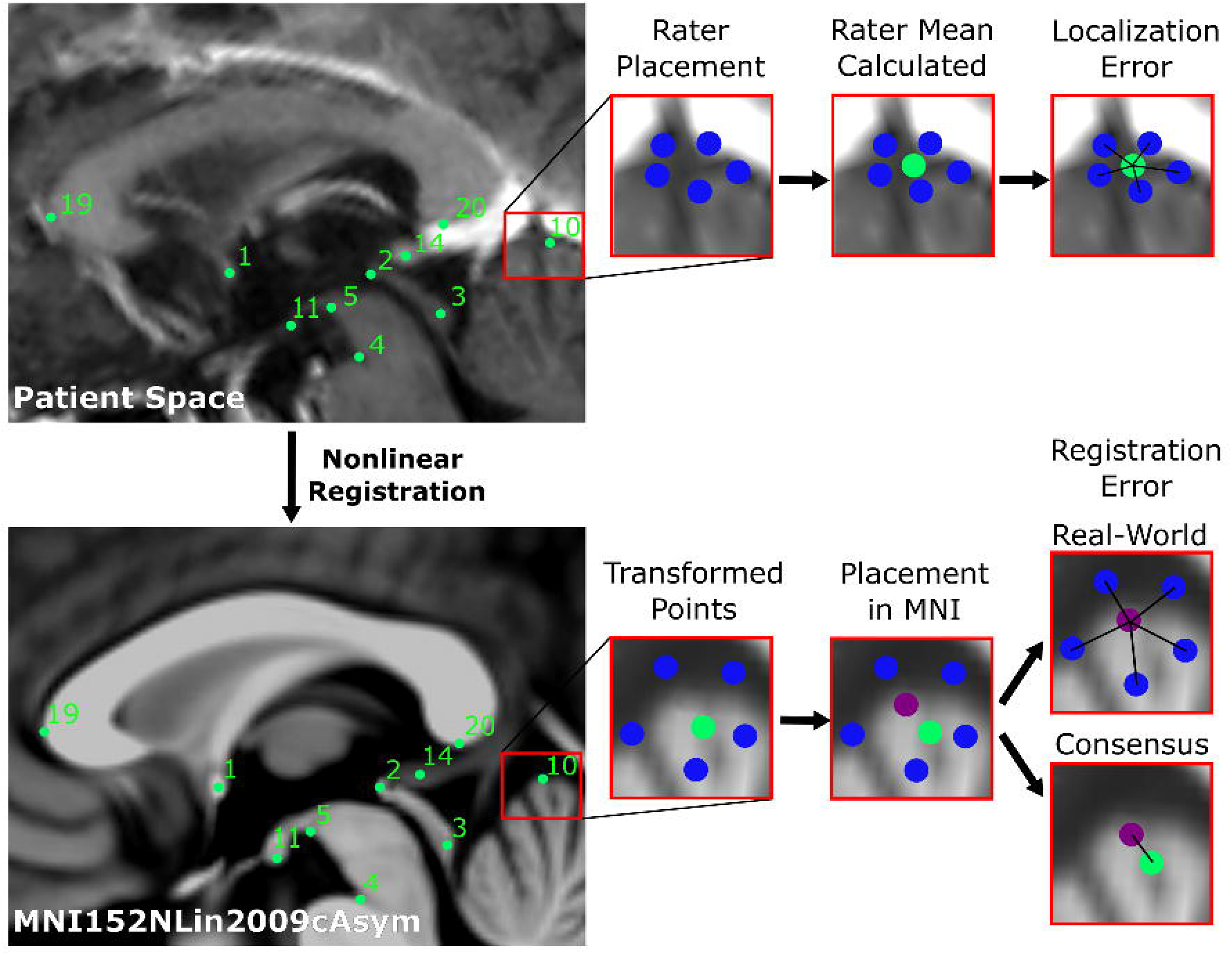
Schematic of workflow to obtain localization errors (above), and registration errors (below). In summary, 5 raters placed 32 anatomical fiducials (AFIDs) on each clinical image (blue). The mean location was calculated for each AFID (green), and the Euclidean distance from each rater’s placement was calculated (termed the localization error). Each rater independently placed AFIDs on the MNI images, and the mean location was calculated (purple). Rater placed AFIDs were transformed to MNI space. The Euclidean distance between each rater’s transformed AFID to the mean location of that AFID placed in MNI space was calculated and termed real-world registration error. Each mean AFID placement on the clinical images was transformed to MNI space, its Euclidean distance to that AFID placed in MNI space was calculated and termed consensus registration error.

We calculated the mean AFRE for linearly and non-linearly registered images. Wilcoxon ranksum tests were used to determine significance between real-world AFREs obtained following both linear and non-linear registration, and significance between non-linearly registered real-world and consensus AFREs. Bonferroni correction was used to account for multiple comparisons with an adjusted p-value of 0.05/32 as a threshold for significance.

### Distance between AFIDs as a biomarker of disease

We sought to investigate a possible secondary benefit of the AFIDs protocol to examine unique morphometric features in our PD patient population. As such, we computed all pairwise Euclidean distances between AFIDs, generating 496 distance measures (32*31/2). We compared these values to distances obtained from a control group of 30 subjects from the OASIS-1 database with AFIDs previously placed (Lau et al. 2019). All 30 subjects used had maximum Mini-Mental State Exam (MMSE) scores (i.e. 30 out of 30). The mean age is 58.0 ± 17.9, and 17 subjects (56.7%) were female. Age between the two groups was compared using an unpaired 2-tailed t-test, and sex between the two groups was compared using a chi-square test. Wilcoxon rank-sum tests were used to determine significant differences in pairwise distances between the two groups, and Bonferroni correction was used to account for multiple comparisons with an adjusted p-value of 0.05/496 being used as a threshold for significance.

## Results

### AFID Placement

Out of all 6240 AFIDs placed, 21 were deemed outliers by using a threshold AFLE of greater than 10 mm (0.33%). None of the outliers were placed by expert raters, therefore 0.55% fiducials placed by novice raters were outliers. All outliers involved placements at some component of the lateral ventricles, and were as follows (number of outliers for this structure in brackets): right lateral ventricle at AC (5), left lateral ventricle at AC (6), right lateral ventricle at PC (2), right anterolateral temporal horn (1), right superior anteromedial horn (1), left superior anteromedial horn (1), right inferior anteromedial horn (2), left inferior anteromedial horn (2) and right ventral occipital horn (1).

### Analysis in Subject Space

Online Resource 2 depicts mean distance from the mid-commissural point by each rater for the 32 AFIDs. The mean overall AFLE across all AFIDs was 1.57 mm ± 1.16 mm. The mean AFLE across all raters for each AFID can be seen in Fig. 2. AFID 25 and 26 (left and right lateral ventricle at AC respectively) had the highest AFLE at 2.63 mm ± 1.75 mm and 2.79 mm ± 1.95 mm respectively. The AFIDs with the lowest overall AFLE were AFID 01-02 (anterior commissure and posterior commissure respectively), with AFLEs of 0.70 mm ± 0.78 mm and 0.55 mm ± 0.34 mm respectively.

**Fig. 2.**
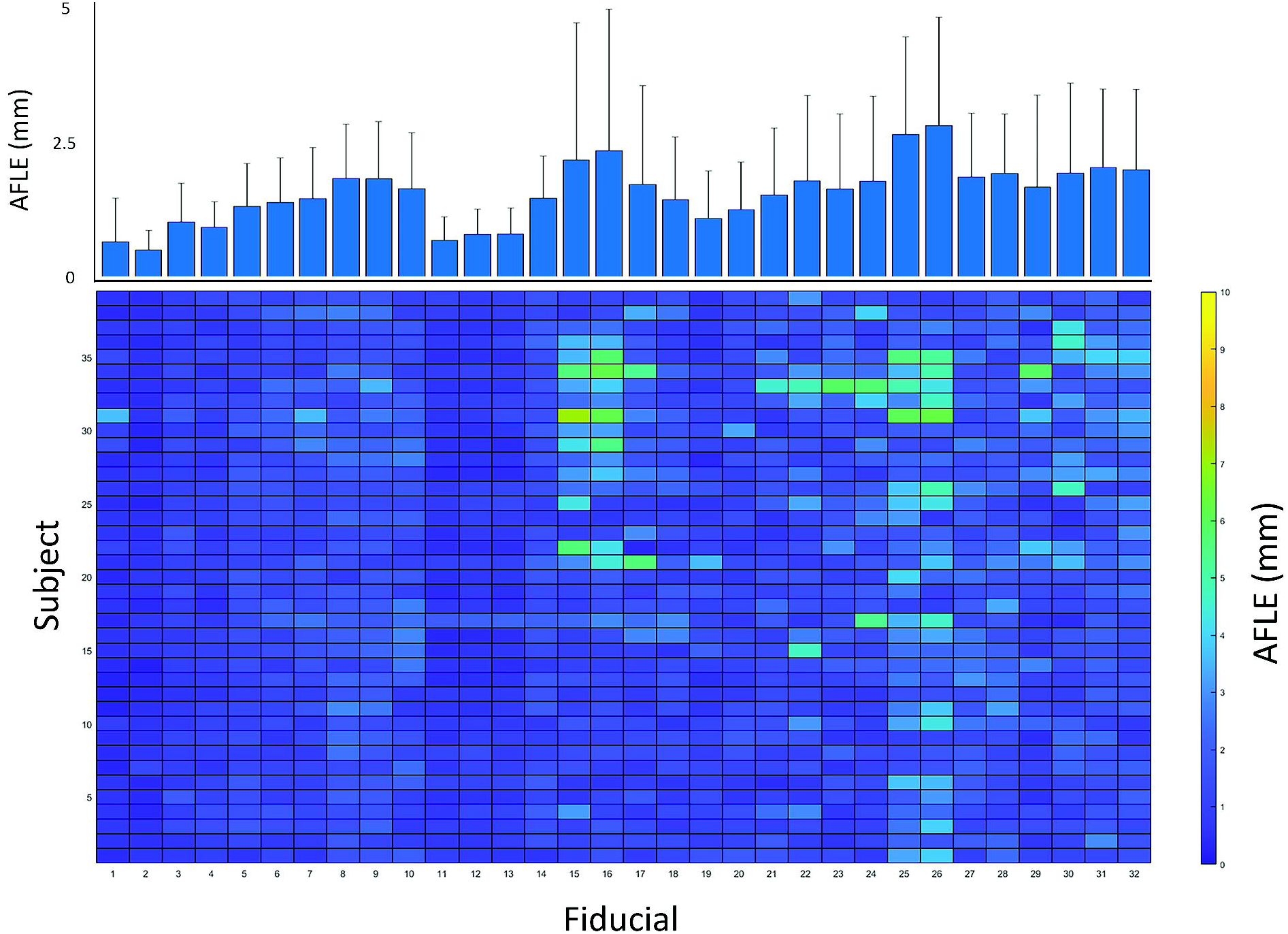
Mean anatomical fiducial localization error (AFLE) for each anatomical fiducial (AFID) and subject. Bottom colormap represents mean AFLEs across all raters for each AFID and subject, illustrating the distribution of AFLEs across all subjects and AFIDs. Top bar graph represents the mean AFLEs for each AFID across all 39 subjects + standard deviation. AFIDs 1, 2 had the lowest AFLEs, while AFIDs 25 and 26 had the greatest AFLEs.

Table 1 represents the mean AFLE obtained by expert and novice raters. Expert raters overall had a lower mean AFLE (1.33 mm ± 0.79 mm), compared to novice raters (1.73 mm ± 1.30 mm). Wilcoxon rank-sum tests for AFLE between expert and novice raters with Bonferroni correction for multiple comparisons are shown in Table 1. Expert raters had a lower AFLE in 29 of the 32 AFIDs. 6 AFIDs had significantly different AFLEs between raters, 5 of which were higher in the novice raters. The superior interpeduncular fossa (AFID05) however had a greater AFLE obtained by expert raters compared to novice raters.

**Table 1.** Mean anatomical fiducial localization error (AFLE) with standard deviation calculated for expert raters (MA and GG) and novice raters (AT, MJ and RC).

| **Fiducial** | **Fiducial Name** | **Expert AFLE (mm)** | **Novice AFLE (mm)** |
| --- | --- | --- | --- |
| 1 | AC | 0.54 ± 0.36 | 0.81 ± 0.45 |
| 2* | PC | 0.41 ± 0.75 | 0.65 ± 1.09 |
| 3 | Infracollicular sulcus | 0.92 ± 1 | 1.15 ± 0.71 |
| 4 | PMJ | 0.86 ± 1.28 | 1.03 ± 1.92 |
| 5* | Superior interpeduncular fossa | 1.60 ± 1.48 | 1.16 ± 1.23 |
| 6 | R superior LMS | 1.35 ± 1.75 | 1.44 ± 1.35 |
| 7 | L superior LMS | 1.55 ± 1.5 | 1.43 ± 1.72 |
| 8 | R inferior LMS | 1.61 ± 1.45 | 1.99 ± 1.9 |
| 9 | L inferior LMS | 1.68 ± 1.27 | 1.94 ± 1.44 |
| 10 | Culmen | 1.35 ± 0.6 | 1.86 ± 0.68 |
| 11 | Intermammillary sulcus | 0.64 ± 0.71 | 0.78 ± 0.85 |
| 12 | R MB | 0.78 ± 0.78 | 0.87 ± 0.92 |
| 13 | L MB | 0.85 ± 1.47 | 0.82 ± 1.34 |
| 14 | Pineal gland | 1.41 ± 1.35 | 1.53 ± 1.29 |
| 15* | R LV at AC | 1.32 ± 1.41 | 2.74 ± 1.45 |
| 16* | L LV at AC | 1.43 ± 1.39 | 2.93 ± 1.2 |
| 17 | R LV at PC | 1.30 ± 1.31 | 2.02 ± 1.02 |
| 18 | L LV at PC | 1.16 ± 1 | 1.65 ± 0.93 |
| 19 | Genu of CC | 0.96 ± 0.89 | 1.22 ± 1.07 |
| 20* | Splenium | 0.98 ± 1.11 | 1.47 ± 1.67 |
| 21 | R AL temporal horn | 1.39 ± 1.35 | 1.64 ± 1.6 |
| 22 | L AL temporal horn | 1.48 ± 1.51 | 2.00 ± 1.38 |
| 23 | R superior AM temporal horn | 1.45 ± 1.56 | 1.78 ± 1.55 |
| 24 | L superior AM temporal horn | 1.56 ± 2.19 | 1.94 ± 2.4 |
| 25 | R inferior AM temporal horn | 2.29 ± 2.57 | 2.85 ± 2.34 |
| 26 | L inferior AM temporal horn | 2.46 ± 1.55 | 3.01 ± 1.48 |
| 27 | R indusium griseum origin | 1.51 ± 1.87 | 2.10 ± 1.64 |
| 28 | L indusium griseum origin | 1.75 ± 1.42 | 2.04 ± 1.26 |
| 29 | R ventral occipital horn | 1.34 ± 1.37 | 1.91 ± 1.58 |
| 30 | L ventral occipital horn | 1.48 ± 2.17 | 2.24 ± 1.3 |
| 31 | R olfactory sulcal fundus | 1.73 ± 1.85 | 2.23 ± 1.25 |
| 32* | L olfactory sulcal fundus | 1.55 ± 1.34 | 2.29 ± 1.33 |
| **Mean** |  | **1.33 ± 0.79** | **1.73 ± 1.30** |

Wilcoxon rank-sum tests were obtained for each anatomical fiducial (AFID) between expert and novice raters, with a significance threshold of 0.05/32. 6 AFIDs had significantly different AFLEs obtained by novice and expert raters. All but one AFID (05) had higher AFLEs obtained by novice raters. AC, anterior commissure; AL, anterolateral; AM, anteromedial; CC, corpus callosum; IPF, interpeduncular fossa; MB, mammillary body; LMS, lateral mesencephalic sulcus; LV, lateral ventricle; PC, posterior commissure; PMJ, pontomesencephalic junction.

To illustrate the differences in AFLE obtained across all 39 subjects and 32 AFIDs, the mean AFLE across the 5 raters was obtained. This produced a 39 by 32 matrix which is represented as a colormap in fig. 2. Each cell in the matrix represents the mean AFLE across the 5 raters for that subject and AFID. This figure illustrates the distribution of errors across the 39 subjects. Some fiducials with a high AFLE, such as AFIDs 25 and 26 demonstrate a consistently higher error across most subjects. However, other fiducials such as AFIDs 15 and 16 only demonstrate a higher AFLE in a subset of subjects.

Intraclass correlation coefficient (ICC) was calculated for each AFID between all raters, expert raters and novice raters, summarized in Table 2. The mean ICC across all AFIDs was 0.814 between all raters, 0.912 between expert raters and 0.777 between novice raters. The superior interpeduncular fossa (AFID 05) had the lowest ICC among both expert and novice raters (0.708 and 0.544 respectively). Otherwise, novice raters also had a lower inter-rater agreement when placing AFIDs associated with the temporal horns (AFIDs 23 to 26). The left anteromedial temporal horn (AFID26) had the second lowest ICC calculated at 0.567 between novice raters, but had an ICC of 0.963 between expert raters.

**Table 2.** Intraclass correlation coefficient (ICC) calculated for each anatomical fiducial (AFID) across 39 subjects, across all raters, expert raters (MA and GG) and novice raters (AT, MJ and RC).

| **Fiducial** | **Fiducial Name** | **Novice ICC** | **Expert ICC** | **Total ICC** |
| --- | --- | --- | --- | --- |
| 1 | AC | 0.674 | 0.958 | 0.771 |
| 2 | PC | 0.855 | 0.964 | 0.895 |
| 3 | Infracollicular sulcus | 0.877 | 0.974 | 0.911 |
| 4 | PMJ | 0.805 | 0.917 | 0.841 |
| 5 | Superior interpeduncular fossa | 0.544 | 0.708 | 0.568 |
| 6 | R superior LMS | 0.726 | 0.822 | 0.747 |
| 7 | L superior LMS | 0.739 | 0.831 | 0.748 |
| 8 | R inferior LMS | 0.796 | 0.885 | 0.814 |
| 9 | L inferior LMS | 0.818 | 0.890 | 0.801 |
| 10 | Culmen | 0.877 | 0.936 | 0.903 |
| 11 | Intermammillary sulcus | 0.798 | 0.826 | 0.816 |
| 12 | R MB | 0.765 | 0.849 | 0.798 |
| 13 | L MB | 0.770 | 0.812 | 0.782 |
| 14 | Pineal gland | 0.756 | 0.835 | 0.757 |
| 15 | R LV at AC | 0.778 | 0.972 | 0.846 |
| 16 | L LV at AC | 0.764 | 0.970 | 0.841 |
| 17 | R LV at PC | 0.762 | 0.967 | 0.830 |
| 18 | L LV at PC | 0.872 | 0.971 | 0.908 |
| 19 | Genu of CC | 0.937 | 0.975 | 0.952 |
| 20 | Splenium | 0.886 | 0.979 | 0.922 |
| 21 | R AL temporal horn | 0.873 | 0.961 | 0.904 |
| 22 | L AL temporal horn | 0.723 | 0.953 | 0.777 |
| 23 | R superior AM temporal horn | 0.706 | 0.876 | 0.755 |
| 24 | L superior AM temporal horn | 0.637 | 0.914 | 0.661 |
| 25 | R inferior AM temporal horn | 0.625 | 0.943 | 0.704 |
| 26 | L inferior AM temporal horn | 0.567 | 0.963 | 0.649 |
| 27 | R indusium griseum origin | 0.829 | 0.931 | 0.866 |
| 28 | L indusium griseum origin | 0.836 | 0.866 | 0.853 |
| 29 | R ventral occipital horn | 0.924 | 0.990 | 0.947 |
| 30 | L ventral occipital horn | 0.926 | 0.991 | 0.946 |
| 31 | R olfactory sulcal fundus | 0.748 | 0.884 | 0.780 |
| 32 | L olfactory sulcal fundus | 0.673 | 0.867 | 0.737 |
| **Mean** |  | **0.777** | **0.912** | **0.814** |

ICC was calculated using a two-way random effects model with a single measurement type. The mean ICC in these three groups was obtained across all AFIDs. AC, anterior commissure; AL, anterolateral; AM, anteromedial; CC, corpus callosum; IPF, interpeduncular fossa; MB, mammillary body; LMS, lateral mesencephalic sulcus; LV, lateral ventricle; PC, posterior commissure; PMJ, pontomesencephalic junction.

### Analysis in MNI Space

To demonstrate the use of AFIDs in determining registration error, subject scans were linearly and non-linearly transformed to the MNI152NLin2009cAsym brain template. The mean real-world and consensus AFREs were calculated. Linear and non-linear real-world AFREs for each AFID are presented in Online Resource 3. The mean non-linear real-world AFRE is 3.34 mm ± 1.94 mm, and the linear AFRE is 4.15 mm ± 2.03 mm. Wilcoxon rank-sum tests for real-world AFREs between linear and non-linear registration with Bonferroni correction for multiple comparisons are shown in Online Resource 3. 15 of the 32 AFIDs had a significantly greater AFRE when using linear registration compared to non-linear registration. Additionally, the consensus AFRE is presented in Online Resource 3, with a mean consensus AFRE of 2.82 mm ± 2.01 mm. 6 AFIDs had a significantly higher non-linear real-world AFRE compared to consensus AFRE (AFIDs 1, 5, 8, 9, 31 and 32).

Fig. 3 demonstrates the mean non-linear real-world AFRE across all subjects and raters for each AFIDs. The anterior commissure (AFID01) had the smallest AFRE calculated at 1.11 mm ± 1.06 mm. The right and left ventral occipital horns (AFIDs 29 and 30) had the largest AFRE at 6.81 mm ± 2.94 mm and 7.36 mm ± 3.41 mm respectively. A colormap of non-linear AFRE across the 5 raters for each subject and AFID is illustrated in fig. 3. This figure demonstrates that the AFIDs with the smallest registration error (AFIDs 1, 2, 11, 12, 13, 31 and 32) were robustly decreased across most subjects. Alternatively, AFIDs 29 and 30 had large registration errors across multiple subjects.

**Fig. 3.**
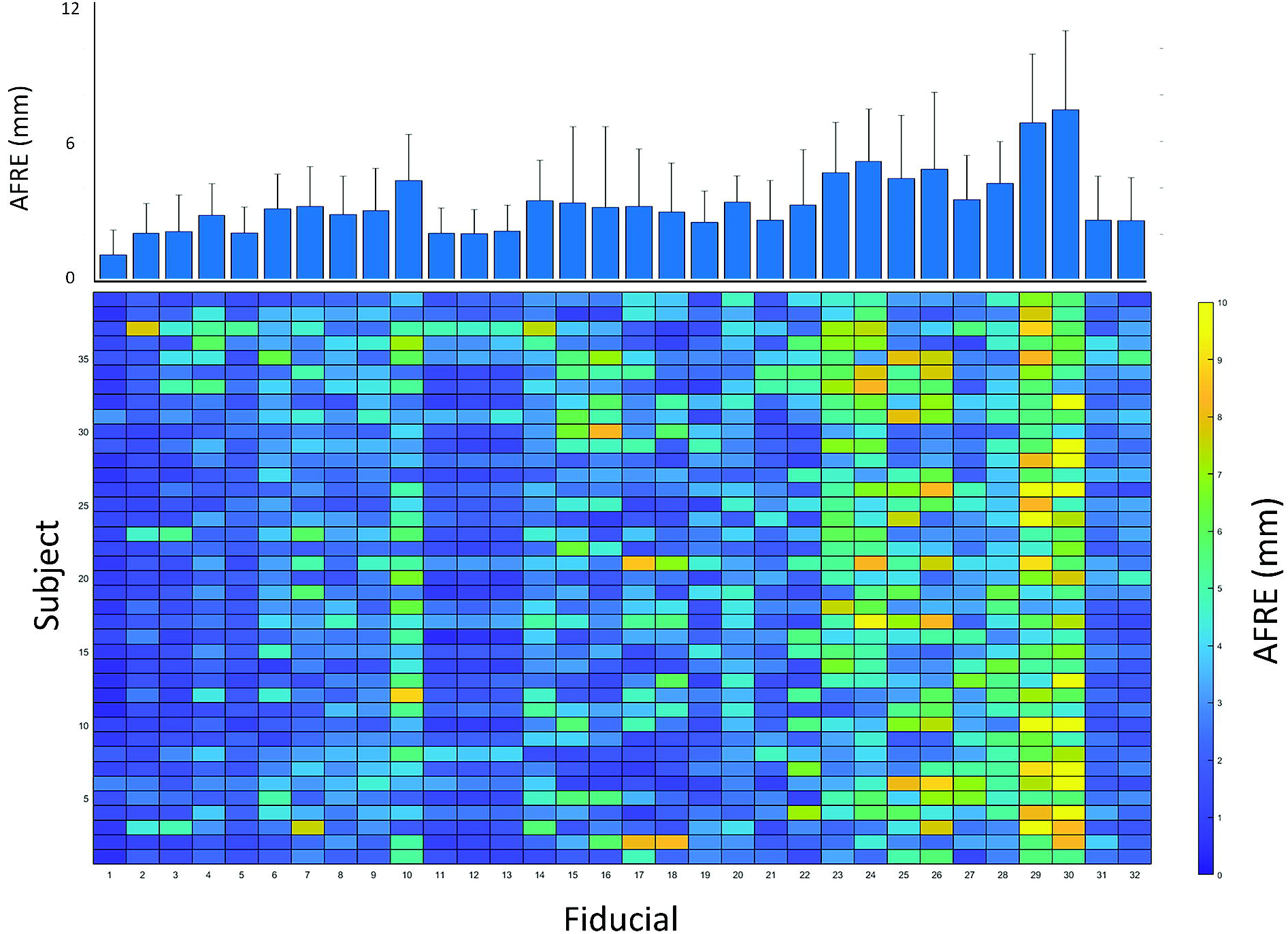
Mean real-world anatomical registration error (AFRE) for each anatomical fiducial (AFID) and subject. Bottom colormap represents mean non-linear AFREs across all raters for each AFID and subject, illustrating the distribution of non-linear AFREs across all subjects and AFIDs. Top bar graph represents the mean non-linear AFRE for each AFID across all 39 subjects + standard deviation. AFIDs 1, 2, 11, 12, 13, 31 and 32 had decreased AFREs across most subjects. AFIDs 29 and 30 had large AFREs across most subjects.

### Distance between AFIDs as a biomarker of disease

An unpaired 2-tailed t-test was performed to compare age across the two groups, demonstrating no statistical difference in age (p = 0.48). Additionally, a chi-square test demonstrated no difference in sex distribution among the two groups (χ2 (1, N=69 = 3.76, p = 0.053).

496 unique Euclidean pairwise distances were calculated between AFIDs for our PD patients (n = 39) and OASIS-1 subjects (n = 30). Fig. 4 represents the differences between the mean of each pairwise distance, calculated by subtracting the mean distances in the OASIS-1 dataset from the mean distances in the PD subject dataset (therefore a positive value indicates a greater pairwise distance in the OASIS-1 subjects). Wilcoxon rank-sum tests were used, and statistically significant differences are indicated in fig. 4. Significance was determined after Bonferroni correction (i.e. by obtaining a p-value < 0.05/496).

**Fig. 4.**
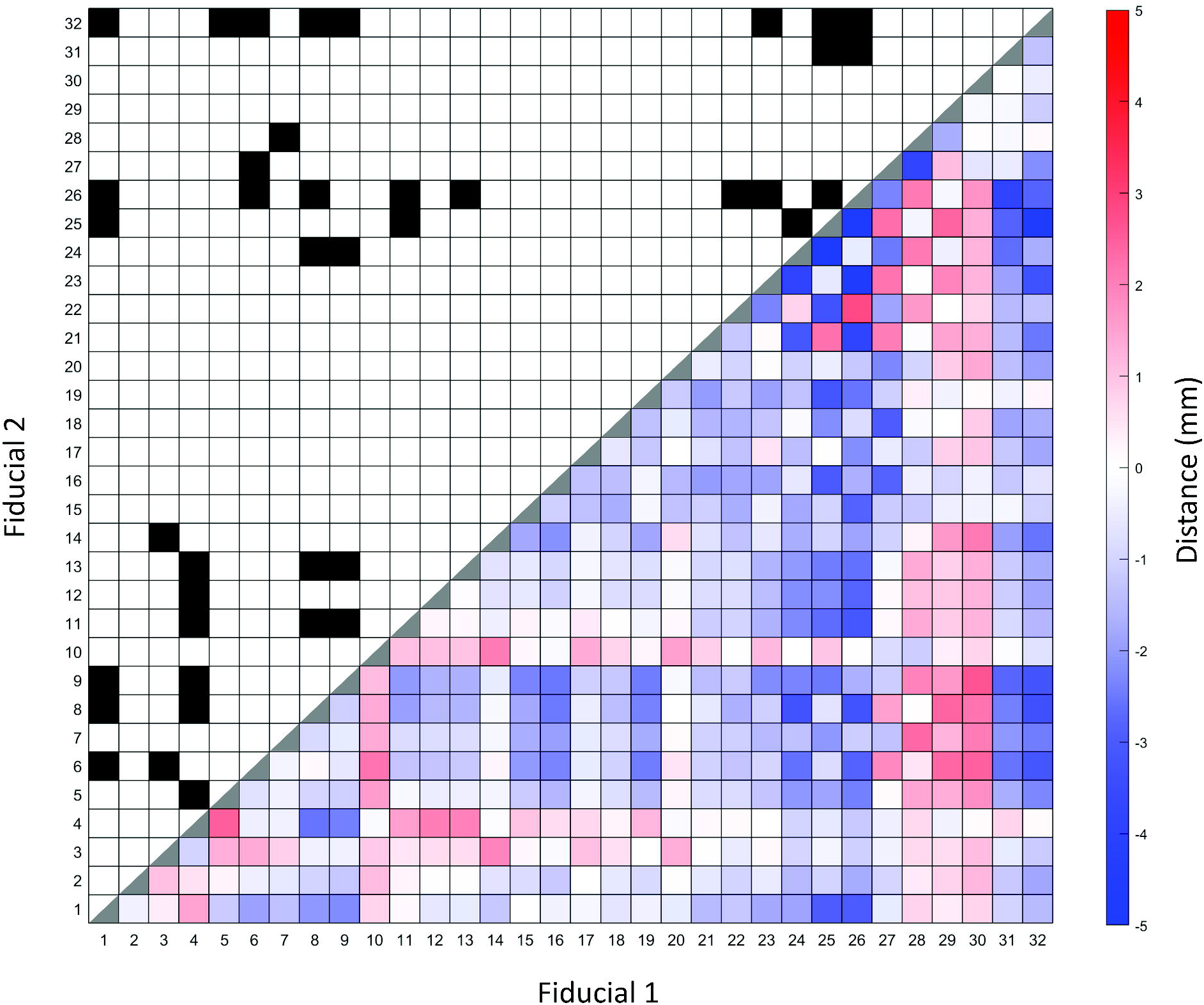
Summary of mean pairwise distances between each anatomical fiducial (AFID) with significant differences. Bottom right shows heatmap representing the difference between mean pairwise distances between each AFID for OASIS-1 subjects and Parkinson’s disease (PD) patients. Positive differences represent a greater pairwise distance in the OASIS-1 subjects relative to PD patients. Significant differences illustrated in top left of figure, designated by a black box. Significance is determined by Wilcoxon rank-sum tests with Bonferroni correction, with a significance threshold of 0.05/496. 40 pairwise distances reached thresholds of statistical significance between PD vs controls (see Online Resource 4 for details).

Between our PD and OASIS-1 datasets, 40 pairwise distances were statistically significantly different (Online Resource 4). The largest distances as a percentage of the distance in the PD dataset were in the left anterolateral to left inferior anteromedial temporal horn (AFID 22 and 26; 2.82 mm, 29.6%), the pontomesencephalic junction to the superior interpeduncular fossa (AFIDs 4 and 5; 2.47 mm, 26.5%) and the infracollicular sulcus to the pineal gland (AFIDs 3 and 14; 1.90 mm, 18.7%).

## Discussion

### Summary

AFIDs were developed as a method to provide a point-based distance to evaluate brain image correspondence (Lau et al. 2019). Using a set of standard templates and individual subject datasets, we previously found that AFID placement protocol was reproducible and more sensitive to local registration error compared to commonly applied voxel overlap measures. The current study sought to apply the AFIDs framework to a clinical dataset, using a set of MRI images obtained in a population of patients with PD. We first demonstrate that AFIDs can be placed with low error by both novices and experts. We then demonstrate the use of AFIDs to evaluate the transformation of our clinical images to a standard MNI brain template. We obtained point-based measures to evaluate local registration error for each subject. Finally, by comparing patients with controls, we provide evidence that the distance between AFIDs could be a biomarker of PD that does not rely on any special imaging scans other than a volumetric structural T1 weighted MRI scan.

### Accuracy of AFID placement

To investigate the accuracy of AFID placement in a clinical setting, we obtained AFLE measurements among novice and expert raters. A mean AFLE of 1.57 mm ± 1.16 mm was obtained across all raters and clinical images. Expert raters generally placed AFIDs with greater accuracy than novice raters, as evidenced by a lower mean AFLE (1.33 mm ± 0.79 mm compared to 1.73 mm ± 1.30 mm) and a greater inter-rater reliability. This suggests that prior knowledge of neuroanatomy does aid in the placement of AFIDs, and that expert raters are more accurate in their use of the AFIDs framework in clinical applications reliant on accurate MRI registration.

The points with lowest AFLE across all raters were the anterior and posterior commissures (AFID01-02; 0.70 mm ± 0.78 mm and 0.55 mm ± 0.34 mm respectively), which has also been found previously (Liu and Dawant 2015; Lau et al. 2019). Points associated with the ventricular system had the largest AFLE. However, we find that errors associated with these AFIDs are not always homogeneously distributed across subjects. For instance, while the right and left inferior anteromedial temporal horns (AFID25-26) had a high AFLE in most subject scans, the right and lateral ventricles at the anterior commissure (AFIDs 15 and 16) only had a high AFLE in select subject scans (Fig. 2). This may be a consequence of the decreased quality clinical images may be subject to, perhaps making some of these structures more difficult to resolve. Anatomical variability across subjects likely also contributes to an increased AFLE.

Overall, our findings suggest that both novice and expert raters are able to place AFIDs within a margin of error in the millimeter range. In fact, our overall AFLE is comparable with errors obtained in our previous work (Lau et al. 2019), with a mean AFLE of 1.27 mm in high resolution template scans and 1.58 mm in individual scans. We had also found that fiducials around the ventricular system had an increased error, with AFLEs in the 2-3 mm range. Therefore, despite the heterogeneous nature of clinically obtained imaging, we provide evidence that AFIDs can be placed with millimetric accuracy.

### Registration error

We evaluated the use of AFIDs to provide a point-based quantitative metric for registration of our clinical images to a standard MNI template. We used fMRIPrep to perform linear and nonlinear techniques for our registration. We found a mean non-linear real-world AFRE of 3.34 mm ± 1.94 mm and mean linear AFRE of 4.15 mm ± 2.03 mm. 15 of the 32 AFIDs had significantly lower AFREs with non-linear registration compared to linear registration. Non-linear registration has evidence for improved registration accuracy; however, accuracy for these registration methods were previously assessed by calculating voxel overlap in specific ROIs (Klein et al. 2009; Modat et al. 2010; Chakravarty et al. 2009). We demonstrated that point-based accuracy measures can provide a more localized quantification of registration error (Lau et al. 2019). In this paper, we provide evidence that the AFIDs protocol can be used in a clinical set of images to provide localized quantification of registration error. Furthermore, we demonstrate a decreased registration errors obtained through non-linear registration compared to linear registration.

Fig. 3 demonstrates the distribution of non-linear registration errors across each AFID and subject. These findings highlight the utility of performing a point-based measure of registration error, as we are able to quantify local areas of registration error for each patient. A schematic such as this may have utility in clinical and research settings where brain image registration is required for a set of subjects, allowing for focal areas of misregistration to be quickly identified. In our particular case, we can see that AFIDs 29 and 30 had a large AFRE across most patients.

We focussed on examining mean real-world AFREs as these values are representative of registration errors obtained in a clinical setting by a single rater. However, as a metric, the real-world AFRE has the disadvantage of representing a combination of both the localization error of a single rater as well as registration error. On the other hand, the consensus AFREs represent the registration error obtained from the mean coordinates in template space, obtained by averaging the coordinates of multiple raters, prior to transformation to MNI space, following the definition from the original manuscript (Lau et al. 2019). Overall, the consensus AFREs are smaller than the real-world AFREs, since the impact of localization error on the measurement is minimized, and represents a more accurate estimation of AFRE although it requires more manual intervention.

Both non-linear real-world and consensus AFREs we obtained were higher than we previously reported using MRI images from the OASIS database (1.80 mm ± 2.09 mm). Registration may have been affected by the variable quality of clinical images, baseline structural differences in PD patients, and the use of gadolinium-enhanced images for which fMRIPrep is not optimized. We elected to use fMRIPrep due to its focus on robustness rather than accuracy, and because it has been demonstrated to achieve accurate registration in the use of traditional voxel overlap measures (Liu and Dawant 2015). We used fMRIPrep in our previous work to define a baseline for future refinement (Lau et al. 2019) and elected to use it in this study to aid in directly comparing our results.

### Distance between AFIDs as a biomarker of disease

AFIDs provided us with the additional opportunity to investigate for potential biomarkers of PD. We compared pairwise distances between AFIDs in our clinical population to control subjects in the OASIS database. A difference in pairwise distance may represent relative morphometric changes (atrophy or hypertrophy) in the cerebral tissue between AFIDs. In our clinical population, we observed the largest differences between the left anterolateral to left inferior anteromedial temporal horn, the pontomesencephalic junction to the superior interpeduncular fossa and the infracollicular sulcus to the pineal gland. These distances were all smaller in our PD patient population.

Voxel-based morphometric studies in PD patients have resulted in inconsistent findings, with conflicting reports of volumetric changes in the substantia nigra (SN) and various cortical areas (Heim et al. 2017; Pyatigorskaya et al. 2014). The SN has reportedly been associated with smaller volumes in patients with PD (Menke et al. 2009; Minati et al. 2007); although other studies have either reported no difference (Péran et al. 2010) or increased volumes in PD patients (Cho et al. 2011). Widespread cortical atrophy has been reported in PD patients with no cognitive impairment (Jubault et al. 2011; Lyoo et al. 2010), and volumetric decrease in the hippocampus and temporoparietal cortex has been associated with cognitive decline in PD patients (Weintraub et al. 2012). Our results may be in keeping with some of these findings. In particular, a decreased distance between the pontomesencephalic junction to the superior interpeduncular fossa and the infracollicular sulcus to the pineal gland may be a manifestation of a decrease in SN volume. Additionally, hippocampal atrophy may result in a decreased distance between the left anteromedial and anterolateral temporal horn. In fact, Camicioli et al. demonstrated a decrease in hippocampal volumes in patients with PD, with an association between decreased left hippocampal volumes and cognitive decline in PD (Camicioli et al. 2003). Finally, these significant differences in local point-wise distance highlight the need to exercise caution when projecting findings in normal controls to patient groups as there can be differences in local brain shape.

### Limitations and future directions

This study has a number of limitations. Although we demonstrate that on average expert raters had a lower AFLE than novice raters, investigating AFLE in 32 AFIDs introduces multiple comparisons which required statistical correction. However, the sample size of this study may not provide sufficient power in demonstrating significant differences in AFLE for each AFID. Additionally, clinical imaging may be subject to variable image quality which can add subjectivity in placing AFIDs. This may result in higher AFLEs and AFREs, although these results may be more representative of the AFIDs framework applied in a clinical setting. Given that PD is a degenerative disease with minimal imaging findings, we are unable to assess the AFIDs framework in a clinical setting with patients who have mass lesions such as brain tumours. Finally, our comparisons of pairwise distances between AFIDs may be confounded by demographic and imaging differences between our PD patient population and the OASIS-1 subjects. Despite this, we provide a novel framework utilizing AFIDs to investigate for biomarkers.

Further work is required to automate the placement of fiducials, providing clinicians with an efficient method to characterize image registration without the subjectivity of manual AFID placement. Although we demonstrate the use of AFIDs to investigate biomarkers for patients with PD, further work is required to further investigate the robustness of our findings and provide more data that can be used to investigate for subtle biomarkers of neurological diseases.

### Conclusion

In summary, we demonstrate that the AFIDs framework can be applied to a clinical population of PD patients with millimetric accuracy. Successful utilization of AFIDs in the context of neurosurgical planning for stereotactic procedures can provide accurate and quantitative measures of image registration, potentially improving outcomes from such procedures. Additionally, we demonstrate how distances between AFIDs could be used as a biomarker to investigate morphological differences in neurodegenerative diseases. AFIDs provide researchers with the benefit of a common, open framework that can be applied across different studies, allowing for an aggregation of clinical datasets and comparisons between various neurological conditions.

## Supporting information

Online Resource 1

Online Resource 2

Online Resource 3

Online Resource 4

## Declarations

### Ethics approval

The study was approved by the Human Subject Research Ethics Board (HSREB) office at the University of Western Ontario (REB# 109045).

### Consent to participate

Not applicable

### Consent for Publication

Not applicable

### Availability of data and material

The datasets generated during and/or analysed during the current study are available at: https://github.com/afids/afids-clinical

### Code availability

Processing scripts used for analysis are available at: https://github.com/afids/afids-clinical

### Authors’ contributions

Study conception and design was performed by Mohamad Abbass, Greydon Gilmore, Terry M. Peters, Ali R. Khan and Jonathan C. Lau. Material preparation and data collection was performed by Mohamad Abbass, Greydon Gilmore, Alaa Taha, Ryan Chevalier and Magdalena Jach. Data analysis and interpretation was performed by Mohamad Abbass, Greydon Gilmore, Terry M. Peters, Ali R. Khan and Jonathan C. Lau. The manuscript was written by Mohamad Abbass, Greydon Gilmore and Jonathan C. Lau. All authors read and approved the final transcript.

## Supplementary Information

**Online Resource 1** - Interactive 3-dimensional brain labelled with all anatomical fiducials.

**Online Resource 2** - Mean individual rater Euclidean distance from mid-commissural point (MCP) for all anatomical fiducials in subject space.

**Online Resource 3** - Mean real-world and consensus anatomical fiducial registration error (AFRE) with standard deviation obtained with linear and non-linear registration of clinical images to MNI space using fMRIPrep.

**Online Resource 4** - List of all mean pairwise distances (mm) ± standard deviation that are significantly different between OASIS-1 subjects and Parkinson’s disease patients.

