## Supplementary material for "Application of the anatomical fiducials framework to a clinical dataset of patients with Parkinson’s disease": Online Resource 1

Clinical AFIDS


Midline Points

Off-Midline Points

1: Anterior Commissure

2: Posterior Commissure

3: Infracollicular Sulcus

4: Pontomesenphalic Junction

5: Superior Interpeduncular fossa

6: R Superior Lateral Mesencepahlic Sulcus

7: L Superior Lateral Mesencepahlic Sulcus

8: R Inferior Lateral Mesencepahlic Sulcus

9: L Inferior Lateral Mesencepahlic Sulcus

10: Culmen

11: Intermammillary Sulcus

12: R Mammillary Body

13: L Mammillary Body

14: Pineal Gland

15: R Lateral Ventricle at Anterior Commissure

16: L Lateral Ventricle at Anterior Commissure

17: R Lateral Ventricle at Posterior Commissure

18: L Lateral Ventricle at Posterior Commissure

19: Genu of Corpus Callosum

20: Splenium

21: R Anterolateral Temporal Horn

22: L Anterolateral Temporal Horn

23: R Superior Anteromedial Temporal Horn

24: L Superior Anteromedial Temporal Horn

25: R Inferior Anteromedial Temporal Horn

26: L Inferior Anteromedial Temporal Horn

27: R Indusium Griseum Origin

28: L Indusium Griseum Origin

29: R Ventral Occipital Horn

30: L Ventral Occipital Horn

31: R Olfactory Sulcal Fundus

32: L Olfactory Sulcal Fundus

view: Left
view: Right
view: Front
view: Back
view: Top
view: Bottom
view: -
