## Supplementary figures and images for "Application of the anatomical fiducials framework to a clinical dataset of patients with Parkinson’s disease"

### Online Resource 2

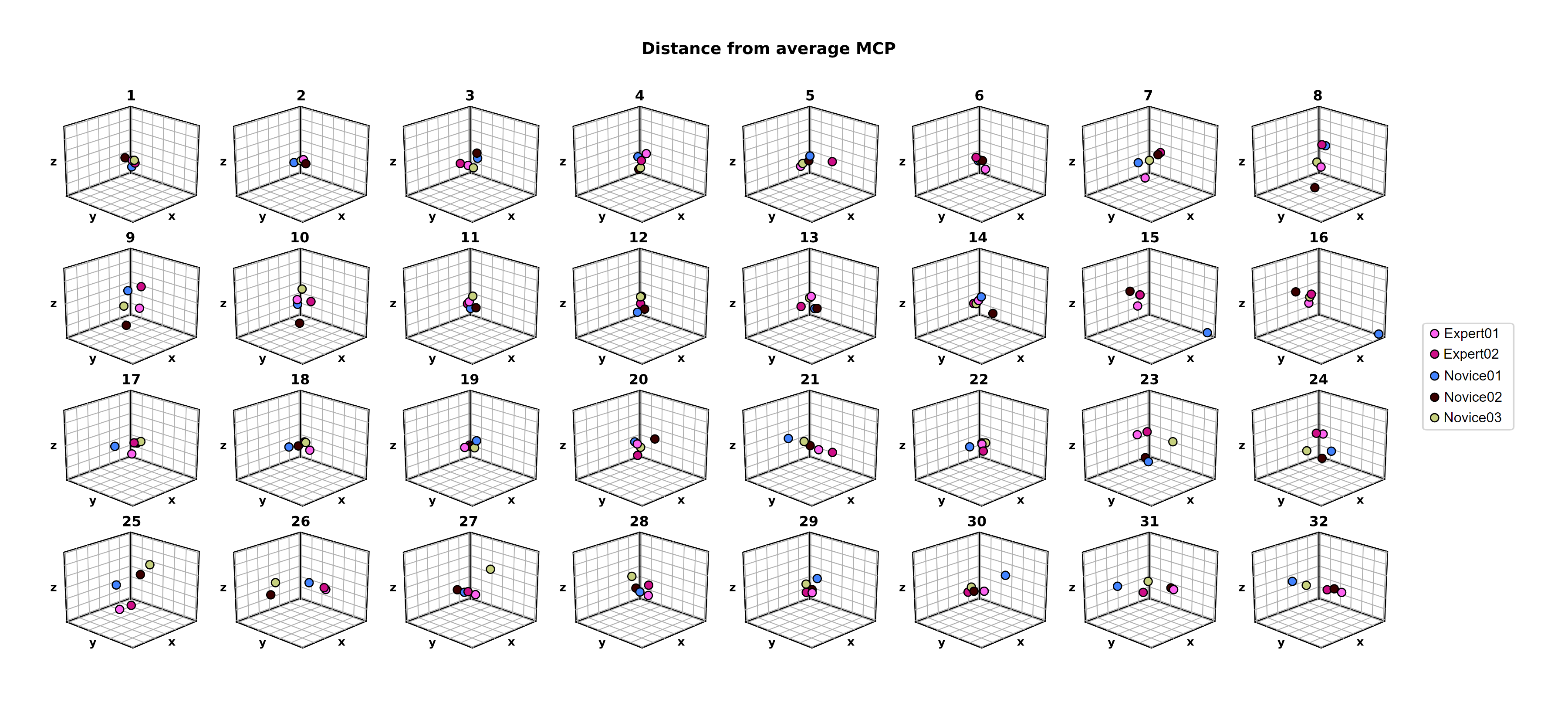
