## Supplementary material for "Application of the anatomical fiducials framework to a clinical dataset of patients with Parkinson’s disease": Online Resource 3

| Fiducial | Fiducial Name | Real-World Linear AFRE (mm) | Real-World Non-linear AFRE (mm) | Consensus Linear AFRE (mm) | Consensus Non-linear AFRE (mm) |
|---|---|---|---|---|---|
| 1*† | AC | 2.39 ± 1.34 | 1.11 ± 1.06 | 2.17 ± 1.20 | 0.78 ± 0.59 |
| 2 | PC | 2.43 ± 1.12 | 2.05 ± 1.27 | 2.34 ± 1.08 | 1.95 ± 1.25 |
| 3 | Infracollicular sulcus | 2.60 ± 1.49 | 2.12 ± 1.56 | 2.30 ± 1.30 | 1.69 ± 1.32 |
| 4* | PMJ | 4.49 ± 1.79 | 2.82 ± 1.36 | 4.40 ± 1.63 | 2.63 ± 1.16 |
| 5*† | Superior interpeduncular fossa | 2.61 ± 1.10 | 2.06 ± 1.12 | 2.12 ± 0.88 | 1.44 ± 0.89 |
| 6 | R superior LMS | 2.80 ± 1.36 | 3.10 ± 1.50 | 2.37 ± 0.85 | 2.66 ± 1.20 |
| 7 | L superior LMS | 2.82 ± 1.43 | 3.21 ± 1.70 | 2.30 ± 0.95 | 2.76 ± 1.40 |
| 8† | R inferior LMS | 3.11 ± 1.54 | 2.86 ± 1.65 | 2.30 ± 1.05 | 1.89 ± 1.04 |
| 9† | L inferior LMS | 3.36 ± 1.54 | 3.01 ± 1.82 | 2.56 ± 1.19 | 2.20 ± 1.15 |
| 10 | Culmen | 3.83 ± 1.74 | 4.32 ± 1.98 | 3.21 ± 1.66 | 3.76 ± 1.74 |
| 11 | Intermammillary sulcus | 2.68 ± 1.51 | 2.05 ± 1.09 | 2.56 ± 1.43 | 1.86 ± 1.03 |
| 12 | R MB | 2.83 ± 1.53 | 2.02 ± 1.05 | 2.68 ± 1.43 | 1.80 ± 0.97 |
| 13 | L MB | 2.89 ± 1.55 | 2.14 ± 1.11 | 2.74 ± 1.45 | 1.95 ± 0.99 |
| 14 | Pineal gland | 2.88 ± 1.33 | 3.45 ± 1.74 | 2.33 ± 1.12 | 3.12 ± 1.28 |
| 15* | R LV at AC | 5.48 ± 3.59 | 3.36 ± 3.28 | 4.93 ± 2.27 | 2.67 ± 1.67 |
| 16* | L LV at AC | 5.71 ± 3.64 | 3.16 ± 3.49 | 4.97 ± 2.54 | 2.51 ± 1.85 |
| 17* | R LV at PC | 4.72 ± 2.71 | 3.21 ± 2.47 | 4.23 ± 2.23 | 2.69 ± 1.91 |
| 18* | L LV at PC | 4.86 ± 2.63 | 2.96 ± 2.10 | 4.49 ± 2.56 | 2.49 ± 1.93 |
| 19* | Genu of CC | 3.81 ± 1.90 | 2.51 ± 1.35 | 3.50 ± 1.86 | 2.25 ± 1.06 |
| 20 | Splenium | 3.17 ± 1.67 | 3.39 ± 1.13 | 2.80 ± 1.46 | 3.25 ± 0.84 |
| 21* | R AL temporal horn | 4.43 ± 1.53 | 2.60 ± 1.72 | 3.89 ± 1.45 | 1.92 ± 1.09 |
| 22* | L AL temporal horn | 5.09 ± 2.00 | 3.26 ± 2.38 | 4.48 ± 1.71 | 2.39 ± 1.74 |
| 23 | R superior AM temporal horn | 4.24 ± 2.27 | 4.66 ± 2.17 | 3.86 ± 1.64 | 4.42 ± 1.53 |
| 24 | L superior AM temporal horn | 5.30 ± 2.67 | 5.14 ± 2.24 | 4.96 ± 1.96 | 4.82 ± 1.63 |
| 25* | R inferior AM temporal horn | 5.94 ± 2.92 | 4.41 ± 2.71 | 5.26 ± 1.96 | 3.27 ± 1.71 |
| 26 | L inferior AM temporal horn | 6.28 ± 3.39 | 4.81 ± 3.30 | 5.70 ± 2.04 | 3.84 ± 2.07 |
| 27* | R indusium griseum origin | 4.50 ± 1.79 | 3.50 ± 1.91 | 3.95 ± 1.47 | 2.82 ± 1.49 |
| 28* | L indusium griseum origin | 5.35 ± 2.15 | 4.20 ± 1.80 | 4.95 ± 1.69 | 3.64 ± 1.33 |
| 29 | R ventral occipital horn | 7.43 ± 2.89 | 6.81 ± 2.94 | 6.99 ± 2.83 | 6.54 ± 2.42 |
| 30 | L ventral occipital horn | 7.42 ± 3.18 | 7.36 ± 3.41 | 6.89 ± 3.25 | 6.86 ± 3.48 |
| 31*† | R olfactory sulcal fundus | 3.52 ± 1.92 | 2.60 ± 1.89 | 2.66 ± 1.35 | 1.77 ± 0.89 |
| 32*† | L olfactory sulcal fundus | 3.84 ± 1.89 | 2.58 ± 1.85 | 3.06 ± 1.37 | 1.58 ± 0.83 |
| **Mean** | | **4.15 ± 2.03** | **3.34 ± 1.94** | **3.69 ± 2.20** | **2.82 ± 2.01** |

Online Resource 3 – Mean real-world and consensus anatomical fiducial registration error (AFRE) with standard deviation obtained with linear and non-linear registration of clinical images to MNI space using fMRIPrep. Wilcoxon rank-sum tests were obtained for each anatomical fiducial (AFID) between linear and non-linear real-world AFRE, with a significance threshold of 0.05/32 (*). Linear AFRE was significantly greater in 15 AFIDs. Wilcoxon rank-sum tests were obtained for each AFID between non-linear real-world and consensus AFRE, with a significance threshold of 0.05/32 (†). Real-world AFRE was significantly greater in 6 AFIDs. AC, anterior commissure; AL, anterolateral; AM, anteromedial; CC, corpus callosum; IPF, interpeduncular fossa; MB, mammillary body; LMS, lateral mesencephalic sulcus; LV, lateral ventricle; PC, posterior commissure; PMJ, pontomesenphalic junction.
