## Supplementary material for "Application of the anatomical fiducials framework to a clinical dataset of patients with Parkinson’s disease": Online Resource 4

| Fiducial 1 | Fiducial 2 | PD Distance | Oasis-1 Distance | Pairwise Difference | Percent Difference |
|---|---|---|---|---|---|
| L AL temporal horn | L inferior AM temporal horn | 9.6 ± 2.5 | 12.4 ± 1.7 | 2.82 | 29.56 |
| PMJ | Superior interpeduncular fossa | 9.3 ± 1.2 | 11.8 ± 1.3 | 2.48 | 26.52 |
| Infracollicular sulcus | Pineal gland | 10.2 ± 1.1 | 12.1 ± 1.2 | 1.90 | 18.70 |
| PMJ | R inferior LMS | 14.8 ± 1.1 | 12.2 ± 1.0 | -2.56 | -17.30 |
| PMJ | L inferior LMS | 14.6 ± 1.0 | 12.2 ± 0.8 | -2.41 | -16.44 |
| PMJ | R MB | 13.4 ± 1.2 | 15.4 ± 1.4 | 2.01 | 15.07 |
| PMJ | L MB | 13.3 ± 1.2 | 15.2 ± 1.3 | 1.98 | 14.96 |
| L superior LMS | L indusium griseum origin | 17.2 ± 1.9 | 19.5 ± 2.3 | 2.33 | 13.52 |
| R inferior AM temporal horn | L inferior AM temporal horn | 46.7 ± 3.7 | 40.7 ± 4.3 | -6.02 | -12.88 |
| PMJ | Intermammillary sulcus | 13.0 ± 1.2 | 14.6 ± 1.4 | 1.51 | 11.56 |
| Intermammillary sulcus | L inferior AM temporal horn | 27.2 ± 2.2 | 24.0 ± 2.7 | -3.12 | -11.48 |
| R superior LMS | R indusium griseum origin | 16.6 ± 1.9 | 18.5 ± 1.7 | 1.86 | 11.16 |
| L superior AM temporal horn | R inferior AM temporal horn | 45.2 ± 3.2 | 40.3 ± 3.6 | -4.90 | -10.84 |
| R superior AM temporal horn | L inferior AM temporal horn | 44.6 ± 3.1 | 39.8 ± 4.0 | -4.74 | -10.63 |
| L inferior LMS | L superior AM temporal horn | 22.3 ± 1.8 | 19.9 ± 2.0 | -2.32 | -10.42 |
| R inferior AM temporal horn | L olfactory sulcal fundus | 44.7 ± 2.5 | 40.1 ± 2.6 | -4.62 | -10.35 |
| L MB | L inferior AM temporal horn | 25.2 ± 2.1 | 22.6 ± 2.4 | -2.60 | -10.31 |
| Intermammillary sulcus | R inferior AM temporal horn | 27.0 ± 2.4 | 24.3 ± 2.5 | -2.69 | -9.98 |
| L inferior AM temporal horn | L olfactory sulcal fundus | 28.5 ± 2.1 | 25.6 ± 2.3 | -2.84 | -9.98 |
| R inferior AM temporal horn | R olfactory sulcal fundus | 29.1 ± 1.8 | 26.3 ± 2.3 | -2.83 | -9.70 |
| AC | L inferior AM temporal horn | 32.7 ± 2.1 | 29.7 ± 2.4 | -3.01 | -9.20 |
| Infracollicular sulcus | R superior LMS | 14.5 ± 0.9 | 15.9 ± 0.9 | 1.34 | 9.20 |
| AC | R inferior AM temporal horn | 32.5 ± 2.3 | 29.6 ± 2.3 | -2.94 | -9.05 |
| R inferior LMS | L superior AM temporal horn | 37.1 ± 2.3 | 33.8 ± 2.5 | -3.27 | -8.82 |
| L inferior AM temporal horn | R olfactory sulcal fundus | 44.5 ± 2.7 | 40.7 ± 3.1 | -3.83 | -8.60 |
| L inferior LMS | Intermammillary sulcus | 25.2 ± 1.5 | 23.2 ± 1.4 | -2.03 | -8.03 |
| R superior AM temporal horn | L olfactory sulcal fundus | 42.6 ± 2.4 | 39.3 ± 2.7 | -3.30 | -7.73 |
| R inferior LMS | Intermammillary sulcus | 25.3 ± 1.6 | 23.3 ± 1.7 | -1.95 | -7.69 |
| R inferior LMS | L inferior AM temporal horn | 42.7 ± 2.6 | 39.4 ± 2.5 | -3.28 | -7.67 |
| AC | L olfactory sulcal fundus | 20.8 ± 1.2 | 19.3 ± 1.3 | -1.47 | -7.10 |
| L inferior LMS | L MB | 24.3 ± 1.5 | 22.7 ± 1.3 | -1.65 | -6.79 |
| Superior interpeduncular fossa | L olfactory sulcal fundus | 34.1 ± 1.8 | 31.8 ± 2.1 | -2.30 | -6.76 |
| L inferior LMS | L olfactory sulcal fundus | 47.6 ± 2.6 | 44.4 ± 2.8 | -3.17 | -6.66 |
| R superior LMS | L inferior AM temporal horn | 44.1 ± 2.8 | 41.3 ± 2.5 | -2.84 | -6.44 |
| R inferior LMS | L olfactory sulcal fundus | 53.2 ± 2.6 | 49.9 ± 2.5 | -3.33 | -6.26 |
| R superior LMS | L olfactory sulcal fundus | 50.2 ± 2.5 | 47.1 ± 2.3 | -3.13 | -6.23 |
| R inferior LMS | L MB | 26.4 ± 1.6 | 24.8 ± 1.6 | -1.59 | -6.03 |
| AC | R superior LMS | 32.0 ± 1.9 | 30.1 ± 1.2 | -1.90 | -5.95 |
| AC | L inferior LMS | 37.3 ± 2.0 | 35.1 ± 1.8 | -2.20 | -5.89 |
| AC | R inferior LMS | 37.4 ± 2.1 | 35.3 ± 1.9 | -2.05 | -5.49 |

Online Resource 4 - List of all mean pairwise distances (mm) ± standard deviation that are significantly different between OASIS-1 subjects and Parkinson's disease patients. Significance is determined by Wilcoxon rank-sum tests with Bonferroni correction, significance threshold of 0.05/496. AC, anterior commissure; AL, anterolateral; AM, anteromedial; CC, corpus callosum; IPF, interpeduncular fossa; MB, mammillary body; LMS, lateral mesencephalic sulcus; LV, lateral ventricle; PC, posterior commissure; PMJ, pontomesenphalic junction.
